# Isoform-specific O-glycosylation dictates Ebola virus infectivity

**DOI:** 10.1101/2022.04.25.489418

**Authors:** Ieva Bagdonaite, Samir Abdurahman, Mattia Mirandola, Martin Frank, Yoshiki Narimatsu, Sergey Y. Vakhrushev, Cristiano Salata, Ali Mirazimi, Hans H. Wandall

## Abstract

Ebola virus glycoprotein is one of the most heavily O-glycosylated viral envelope glycoproteins. Using glycoengineered cell lines we demonstrate that O-linked glycan truncation and perturbed initiation retarded the production of viral particles and decreased infectivity of progeny virus. Next, using TMT isobaric labelling, we performed quantitative differential O-glycoproteomics on proteins produced in wild type HEK293 cells and cell lines ablated for three key GalNAc-transferases, GalNAc-T1, -T2, and -T3, as well as compared it to patterns on wild type virus-like particles. We demonstrate selective initiation of a subset of O-glycosites by each enzyme, with GalNAc-T1 having the largest contribution. This work represents a comprehensive site-specific analysis of EBOV GP, with 47 O-glycosites identified, and sheds light on differential regulation of EBOV GP glycosylation initiated by host GalNAc-Ts. Together with the effect on viral propagation it opens prospective avenues for tailored intervention approaches and means for modulating immunogen O-glycan density.

## Introduction

Ebola virus (EBOV) is an enveloped filovirus that causes recurrent outbreaks of deadly hemorrhagic fever in Africa. The zoonotic nature of the virus poses a formidable threat in densely populated regions. The most recent outbreaks in West Africa (EBOV Makona) and The Democratic Republic of Congo in 2014 and 2018-2020, respectively, were the most widespread so far and claimed numerous lives. This also led to a search of means to tame the disease and resulted in development of therapeutic antibody cocktails and vaccines (Audet et al., 2014; Bornholdt et al., 2016; Gilchuk et al., 2020; Matz et al., 2019; Regules et al., 2017). However, affordable and widely available treatments are still lacking.

The EBOV glycoprotein (GP) is the only viral protein on the surface of the virion and is critical for attachment to host cells and catalysis of membrane fusion. The GP is a transcriptionally edited transmembrane glycoprotein which is proteolytically processed by a furin-like host protease to yield covalently linked GP1 and GP2 that form trimers of heterodimers (Fig. 1A, 1B) (Volchkov et al., 1998). GP1 mediates cellular attachment in a macropinocytosis-dependent manner by engaging TIM-1, DC-SIGN and other C-type lectins, whereas GP2 executes fusion upon interaction with NPC intracellular cholesterol transporter 1 (NPC1) that thereby acts as an endosomal entry receptor (Gong et al., 2016; Lee et al., 2017; Nanbo et al., 2010; Wang et al., 2016; Zhang et al., 2022). A substantial portion of GP1 is comprised of heavily glycosylated glycan cap domain (GCD) and mucin-like domain (MLD), which collectively occlude the NPC1 binding site, and therefore need to be cleaved off by cathepsins L and B for proper infectivity (Fig. 1A, 1B). Furthermore, an additional priming event is needed in the endosomal compartment to ensure interaction with NPC1 and fusion (Bale et al., 2011; Chandran et al., 2005; Hood et al., 2010; Kaletsky et al., 2007). Glycosylation of viral envelope glycoproteins rely on the glycosylation machinery of the host (Bagdonaite and Wandall, 2018). The most predominant types of glycans in humans include N-linked glycans and mucin-type O-linked glycans (Schjoldager et al., 2020). N-linked glycans modify asparagine residues within N-X-S/T (X ≠ P) sequons and have extensively been characterized on various viral envelope glycoproteins and are easy to predict (Bagdonaite et al., 2018; Shajahan et al., 2020; Watanabe et al., 2020; Zhao et al., 2020). O-glycan initiation is regulated by the competing action of 20 isoforms of polypeptide GalNAc-transferases in humans (Bennett et al., 2012). While there is some knowledge on isoform-specific protein substrates, their non-redundant roles in glycosylating viral envelope proteins have not been addressed (Dabelsteen et al., 2020; Lavrsen et al., 2018; Narimatsu et al., 2019a; Schjoldager et al., 2012). Up to eight core structures have been described for O-GalNAc glycans, though core 1 and core 2 are the most ubiquitously encountered (Fig. 1C) (de Haan et al., 2022; Wandall et al., 2021). GP1 contains 17 N-glycosylation sequons in both the GCD and the MLD; 12 of them are predicted to be utilized (Fig. 1A). A few specific N-glycosites affecting EBOV conformational stability or immunogenicity have been identified and shown to provide steric shielding of host cell ligands for immune effector cells (Dowling et al., 2007; Iraqi et al., 2020). The MLD is also predicted to be heavily glycosylated, though no specific sites have been identified, and the studies on O-glycosylation have been limited to deletion of MLD (Dowling *et al*., 2007; Feldmann et al., 1994; Lee and Saphire, 2009). The extent and role of O-glycosylation in EBOV biology is thus still obscure. Moreover, the structure of the MLD, comprising at least half of GP’s mass (Fujihira et al., 2018), is poorly defined, as the region is usually omitted from crystallographic studies (Fig. 1B) (Rutten et al., 2020).

**Figure 1.**
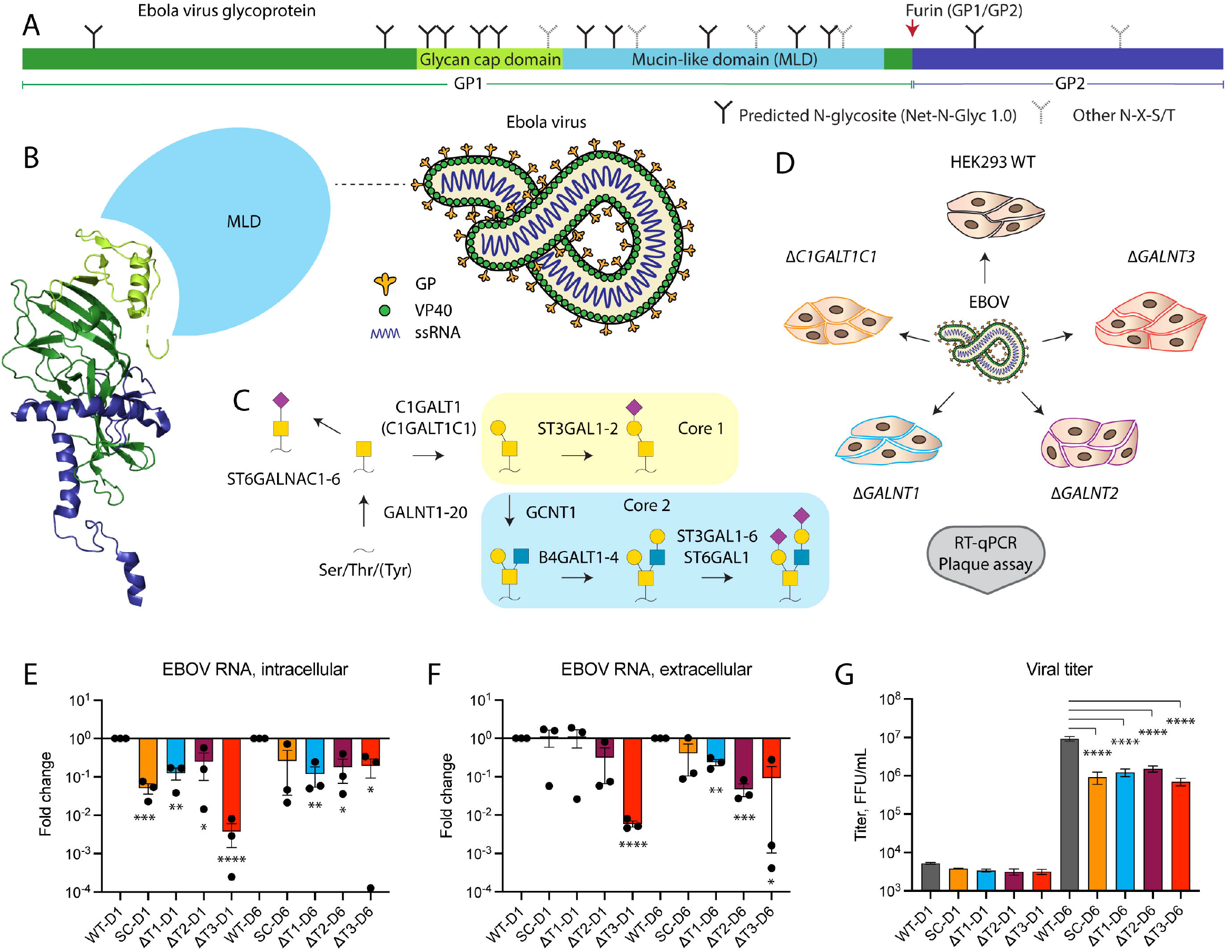
O-glycosylation is important for EBOV infectivity. (A) Layout of Ebola virus glycoprotein. (B) Cartoon depiction of Ebola virus, as well as a ribbon diagram of a monomeric viral envelope glycoprotein (PDB: 6VKM). Color coding as in (A). The mucin-like domain is not resolved and shown as a light blue sphere. (C) Predominant O-glycosylation pathways in HEK293 cells. (D) Experimental strategy for addressing the functional role of O-glycosylation in EBOV biology. (E, F) RT-PCR analysis of viral RNA day 1 (D1) and day 6 (D6) post-infection of glycoengineered HEK293 cells. Expression levels are normalized to β-actin and presented as fold change compared to wild type. Data is shown as mean ±SEM of three independent experiments, where individual datapoints are also shown. One sample t test was used to evaluate differences from 1 (* *p* < 0.05, ** *p* < 0.01, *** *p* < 0.001, **** *p* < 0.0001). (E) Intracellular RNA. (F) Extracellular RNA. (G) Plaque titration analysis of Ebola virus in cell culture supernatants of infected glycoengineered cells on day 1 (D1) and day 6 (D6) post-infection. Data is shown as mean ±SEM of two independent experiments. One-way ANOVA followed by Šidák’s multiple comparison test was used to evaluate differences in titers compared to wild type on the individual days post-infection (* *p* < 0.05, ** *p* < 0.01, *** *p* < 0.001, **** *p* < 0.0001).

Mutational studies have identified the MLD being responsible for EBOV-induced cytotoxicity and vascular permeability (Yang et al., 2000). MLD has also been hypothesized to protect from antibody neutralization, as removal of glycan cap during cathepsin processing would expose immunodominant residues important for receptor binding or cell fusion, and better neutralization of *in vitro* pre-processed pseudotyped particles by convalescent plasma was achieved (Luczkowiak et al., 2018). Given the involvement of the MLD in various stages of viral life cycle and immune recognition, it is important to better understand its structural features, including the predominant glycan structures, their sites, and the regulation of glycan density by GalNAc-transferases. Here, we used tandem mass spectrometry to map O-glycosylation sites on Makona strain EBOV GP and determine the individual contribution of GalNAc-T1, -T2, and -T3 to O-glycan initiation. Furthermore, we investigated the role of O-linked glycan initiation and elongation for EBOV propagation using genetically engineered cell lines and discovered requirement of all five investigated gene products for proper infectivity of progeny virus.

## Results

### EBOV exhibits diminished propagation in O-glycosylation mutant cell lines

In order to address the function of O-glycans in EBOV biology, we infected HEK293 cells lacking O-linked glycan elongation (*C1GALT1C1* KO), or deficient in individual isoforms of predominant initiating enzymes (*GALNT1* KO, *GALNT2* KO or *GALNT3* KO) (Fig. 1D). Intracellular viral replication was diminished in all mutant cell lines compared to wild type on both Day 1 and Day 6 post infection (Fig. 1E). The trend was also reflected in amounts of extracellular viral RNA on Day 6 post infection, with deletion of individual *GALNT* genes having a larger effect compared to elimination of O-glycan elongation (Fig. 1F). We then measured viral titers in cell culture supernatants to evaluate the infectivity of extracellular virions. On Day 1 post-infection, viral titers in supernatants of all four knock out cell lines were slightly lower but still comparable to wild type (Fig. 1G). In contrast, a substantial reduction in viral titers was observed in all four knock out cell lines on Day 6 post-infection (Fig. 1G). The combined data suggests the importance of both O-glycan initiation by the individual GalNAc-T isoforms and O-glycan elongation for sustained viral propagation.

### EBOV GP is expressed on the surface of glycoengineered cells

Reduced titers of extracellular virions could result from a number of different issues, including but not limited to viral envelope GP expression, trafficking, and function. To assess EBOV GP expression in glycogene knock out cell lines (*C1GALT1C1* KO, *GALNT1* KO, *GALNT2* KO and *GALNT3* KO (Fig. S1)) we investigated both native virus-infected cells (Fig. 2A, 2B), and those transfected with a recombinant full length GP (Fig. 2C). Immunofluorescence staining of infected wild type cells at low MOI revealed smaller and larger clusters of infected cells with predominant surface expression of GP (Fig. 2A, 2B). Aside from *GALNT2* KO cells often exhibiting very small lesions (Fig. 2A, 2B *GALNT2* KO lower images), all KO cell lines appeared competent for GP surface expression. To confirm GP surface expression in non-permeabilized cells, we expressed full length EBOV GP in the suite of glycoengineered cell lines and co-stained them with surface marker E-cadherin (Fig. 2C). In agreement with infected cell data, recombinant GP was detected on the surface of all five cell lines. This suggests that alterations in O-glycosylation do not impair glycoprotein trafficking. Furthermore, it allowed us to use the glycoengineered cell lines for recombinant GP production for the analysis of how the individual GalNAc-T isoforms define the O-glycosylation pattern of GP.

**Figure 2.**
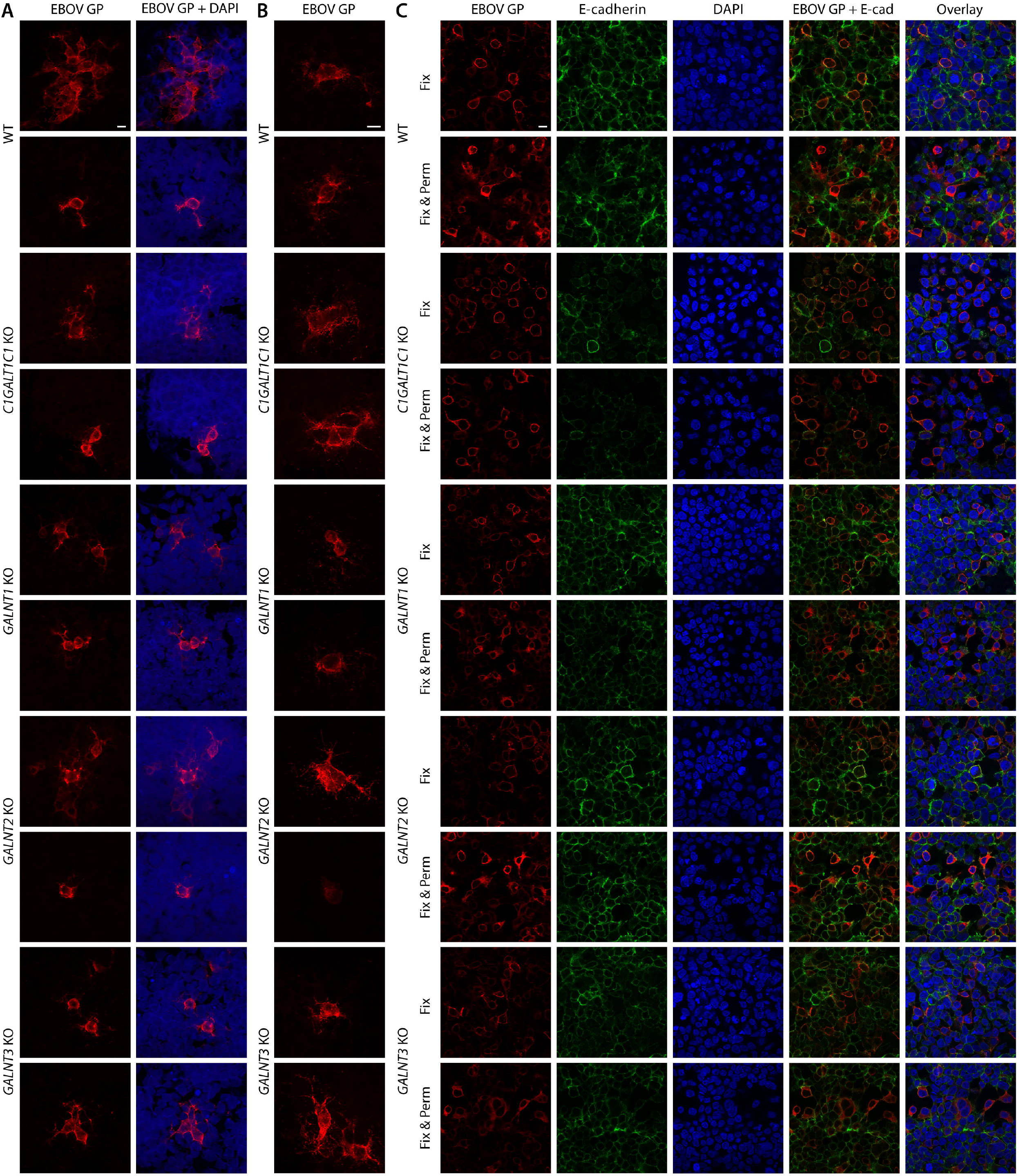
EBOV GP expression in glycoengineered cell lines. (A, B) HEK293 knock out cell lines infected with the Makona isolate of Zaire EBOV at 0.01 MOI and fixed in acetone 24 hours post-infection were stained for EBOV GP (red). Two images for each cell line are shown. (A) Confocal snapshots. Scale bar 10 μm. (B) z-stack maximal intensity projections. Scale bar 10 μm. (C) Indicated cell lines transfected with a plasmid encoding full length EBOV Makona GP were fixed with 4 % PFA (Fix) 48 hours post-tranfection and co-stained for GP (red) and E-cadherin (green). Another set of cells was also permeabilized with 0.3 % Triton X-100 (Fix & Perm). Scale bar 10 μm.

### Different GalNAc-T isoforms glycosylate distinct sites within the MLD

Cumulative reduction of viral titers over time suggested that modulation of O-glycan length and density may have an effect on MLD stability or function. We therefore wanted to generate a more comprehensive knowledge of the O-glycans that decorate the MLD and next mapped specific O-glycosites, their predominant structures, and site-selective initiation by GalNAc-T isoforms. In order to identify O-linked glycosylation sites on EBOV GP, we analyzed EBOV virus-like particles (VLPs) presenting the GP on native-like rod-shaped particles generated by co-expression with VP40 protein representing the mature glycoprotein (Fig. 3A). To determine the individual contributions of GalNAc-T1, -T2, and T3, we expressed the full length His-tagged EBOV GP in WT and the different *GALNT* KO cell lines, purified and digested the protein, and labelled the resulting peptides with TMT isobaric mass tags allowing for relative quantification of (glyco)peptides in the different samples (Fig. 3B) (Hintze et al., 2018; Mao et al., 2021). In both approaches, we performed sequential lectin weak affinity chromatography (LWAC) of desialylated tryptic digests to enrich for abundant core 1 O-glycan structures (T, Galβ1-3GalNAcα1-O-Ser/Thr) and their biosynthetic intermediates (Tn, GalNAcα1-O-Ser/Thr) using peanut agglutinin (PNA) and *Vicia villosa* lectin (VVA), respectively. We identified 47 O-glycosites, 45 of which unambiguous, majority of which located to MLD and glycan cap regions and were in good agreement between VLPs and purified GP, with 32 O-glycosites in common (Fig. 4A, Table 1, Dataset S1). PNA LWAC allowed for enrichment of not only core 1, but also core 2 structures, which were abundantly found at the majority of glycosites. Notably, core 2 structures could co-exist on adjacent amino acids, such as Thr326 and Ser327, Ser347 and Ser348, or Thr424 and Thr425. In contrast, we found a few GP stretches, where no elongation was taking place due to dense O-glycosylation, with up to seven glycosites identified on the same peptide (366-380), predominantly as single GalNAc structures. This signifies local structural differences in glycosyltransferase accessibility within different subregions of the MLD.

**Table 1.** O-glycosites and structures.

| Glycosite | Recombinant GP PNA | Recombinant GP FT VVA | GalNAc-T regulation | Median KO/WT | VLPs PNA | VLPs FT VVA |
| --- | --- | --- | --- | --- | --- | --- |
| Thr42 |  |  |  |  | Hex1 HexNAc1 | HexNAc |
| Thr77 |  | HexNAc | GalNAc-T1 | 0.43 |  |  |
| Thr83 |  | HexNAc |  |  |  |  |
| Ser195 |  |  |  |  | Hex1 HexNAc1 |  |
| Thr206 |  | HexNAc | GalNAc-T1;<br>GalNAc-T2 | 0.42;<br>0.27 | Hex1 HexNAc1,<br>Hex2 HexNAc2 | HexNAc |
| 283-284 |  |  |  |  | Hex1 HexNAc1 |  |
| Thr309 | Hex1HexNAc1,<br>Hex1HexNAc2,<br>Hex2HexNAc2 |  |  |  | Hex1 HexNAc1,<br>Hex2 HexNAc2 | HexNAc |
| Ser312 | HexNAc,<br>Hex1HexNAc1,<br>Hex1HexNAc2,<br>Hex2HexNAc2 | HexNAc | GalNAc-T1;<br>GalNAc-T3 | 0.31;<br>0.33 | Hex1 HexNAc1,<br>Hex2 HexNAc2 | HexNAc |
| Ser319 |  |  |  |  | HexNAc |  |
| Ser322 |  |  |  |  | HexNAc,<br>Hex1 HexNAc1 |  |
| Thr326 | Hex1HexNAc1,<br>Hex1HexNAc2,<br>Hex2HexNAc2 | HexNAc |  |  | HexNAc<br>Hex1 HexNAc1,<br>Hex2 HexNAc2 |  |
| Ser327 | HexNAc,<br>Hex1HexNAc1,<br>Hex1HexNAc2,<br>Hex2HexNAc2 |  | GalNAc-T2 | 0.09 | HexNAc,<br>Hex1 HexNAc1,<br>Hex1 HexNAc2,<br>Hex2 HexNAc2 | HexNAc |
| Ser328 |  | HexNAc |  |  | HexNAc,<br>Hex1 HexNAc1,<br>Hex2 HexNAc2 |  |
| Thr332 |  | HexNAc |  |  | HexNAc,<br>Hex1 HexNAc1,<br>Hex2 HexNAc2 |  |
| Thr334 | Hex1HexNAc1,<br>Hex1HexNAc2,<br>Hex2HexNAc2 | HexNAc | GalNAc-T1 | 0.19 | Hex2 HexNAc2 |  |
| Thr335 |  |  |  |  | Hex1 HexNAc2 | HexNAc |
| Ser344 | HexNAc,<br>Hex2HexNAc2 |  |  |  | HexNAc,<br>Hex1 HexNAc1,<br>Hex2 HexNAc2 |  |
| Ser347 | HexNAc,<br>Hex1HexNAc1,<br>Hex2HexNAc2 |  |  |  | HexNAc,<br>Hex2 HexNAc2 |  |
| Ser348 | HexNAc,<br>Hex1HexNAc1,<br>Hex1HexNAc2,<br>Hex2HexNAc2 | HexNAc,<br>Hex1HexNAc1,<br>Hex2HexNAc2 | GalNAc-T1 | 0.22 | HexNAc,<br>Hex1 HexNAc1,<br>Hex2 HexNAc2 | HexNAc |
| Thr355 | Hex2HexNAc2 | HexNAc |  |  | HexNAc,<br>Hex1 HexNAc1,<br>Hex2 HexNAc2 |  |
| Thr366 | HexNAc | HexNAc |  |  |  | HexNAc |
| Thr367 | HexNAc | HexNAc |  |  | Hex1HexNAc1<br>(1x) | HexNAc |
| Thr370 | HexNAc (2x) | HexNAc |  |  |  | HexNAc |
| Ser372 |  |  |  |  | Hex1HexNAc1<br>(1x) | (1x) |
| Thr373 |  | HexNAc |  |  |  | HexNAc |
| Ser374 | Hex1HexNAc1 | HexNAc |  |  |  | HexNAc |
| Thr379 | HexNAc | HexNAc |  |  | Hex1HexNAc1 | HexNAc |
| Thr380 | HexNAc | HexNAc |  |  |  | HexNAc |
| Thr382 |  | HexNAc |  |  |  | HexNAc |
| Thr388 | Hex1HexNAc1,<br>Hex2HexNAc2 | HexNAc | GalNAc-T1 | 0.47 | HexNAc | HexNAc |
| Thr391 | Hex1HexNAc1 | HexNAc |  |  | Hex1HexNAc1,<br>Hex1HexNAc2,<br>Hex2HexNAc2 | HexNAc |
| Ser399 | HexNAc,<br>Hex1HexNAc1,<br>Hex1HexNAc2,<br>Hex2HexNAc2 | HexNAc |  |  | HexNAc,<br>Hex1HexNAc1,<br>Hex1HexNAc2,<br>Hex2HexNAc2 | HexNAc |
| Thr402 | HexNAc,<br>Hex1HexNAc1,<br>Hex1HexNAc2,<br>Hex2HexNAc2 | HexNAc | GalNAc-T1 | 0.43 | HexNAc,<br>Hex1HexNAc1,<br>Hex1HexNAc2,<br>Hex2HexNAc2 | HexNAc |
| Thr416 | HexNAc | HexNAc |  |  | HexNAc<br>Hex1HexNAc1 | HexNAc |
| Ser418 | HexNAc,<br>Hex1HexNAc2 | HexNAc | GalNAc-T1 | 0.31 | HexNAc,<br>Hex1HexNAc1 |  |
| Thr420 | Hex1HexNAc1 | HexNAc |  |  |  |  |
| Thr424 | HexNAc,<br>Hex1HexNAc1,<br>Hex1HexNAc2,<br>Hex2HexNAc2 | HexNAc |  |  | Hex1HexNAc1,<br>Hex2HexNAc2 | HexNAc |
| Thr425 | HexNAc,<br>Hex1HexNAc1,<br>Hex1HexNAc2,<br>Hex2HexNAc2 | HexNAc |  |  | HexNAc,<br>Hex1HexNAc1,<br>Hex2HexNAc2 | HexNAc |
| Thr435 |  | HexNAc |  |  |  |  |
| Ser438 | HexNAc,<br>Hex1HexNAc1,<br>Hex1HexNAc2,<br>Hex2HexNAc2 | HexNAc | GalNAc-T1 | 0.13 | HexNAc,<br>Hex1HexNAc1 | HexNAc |
| Thr448 | Hex1HexNAc1 | HexNAc |  |  |  |  |
| Thr449 |  | HexNAc |  |  |  |  |
| Thr450 |  | HexNAc |  |  |  |  |
| 455-456<br>(1x) | Hex1HexNAc1 |  |  |  |  |  |
| Ser475 |  | HexNAc | GalNAc-T1 | 0.06 |  |  |
| Thr483 |  | HexNAc |  |  | Hex1HexNAc1,<br>Hex2HexNAc2 |  |
| Thr485 | Hex2HexNAc2 | HexNAc |  |  | Hex1HexNAc1,<br>Hex2HexNAc2 | HexNAc |
| Thr494 | Hex1HexNAc1,<br>Hex2HexNAc2 | HexNAc | GalNAc-T3 | 0.3 | Hex1HexNAc1,<br>Hex2HexNAc2 | HexNAc |

**Figure 3.**
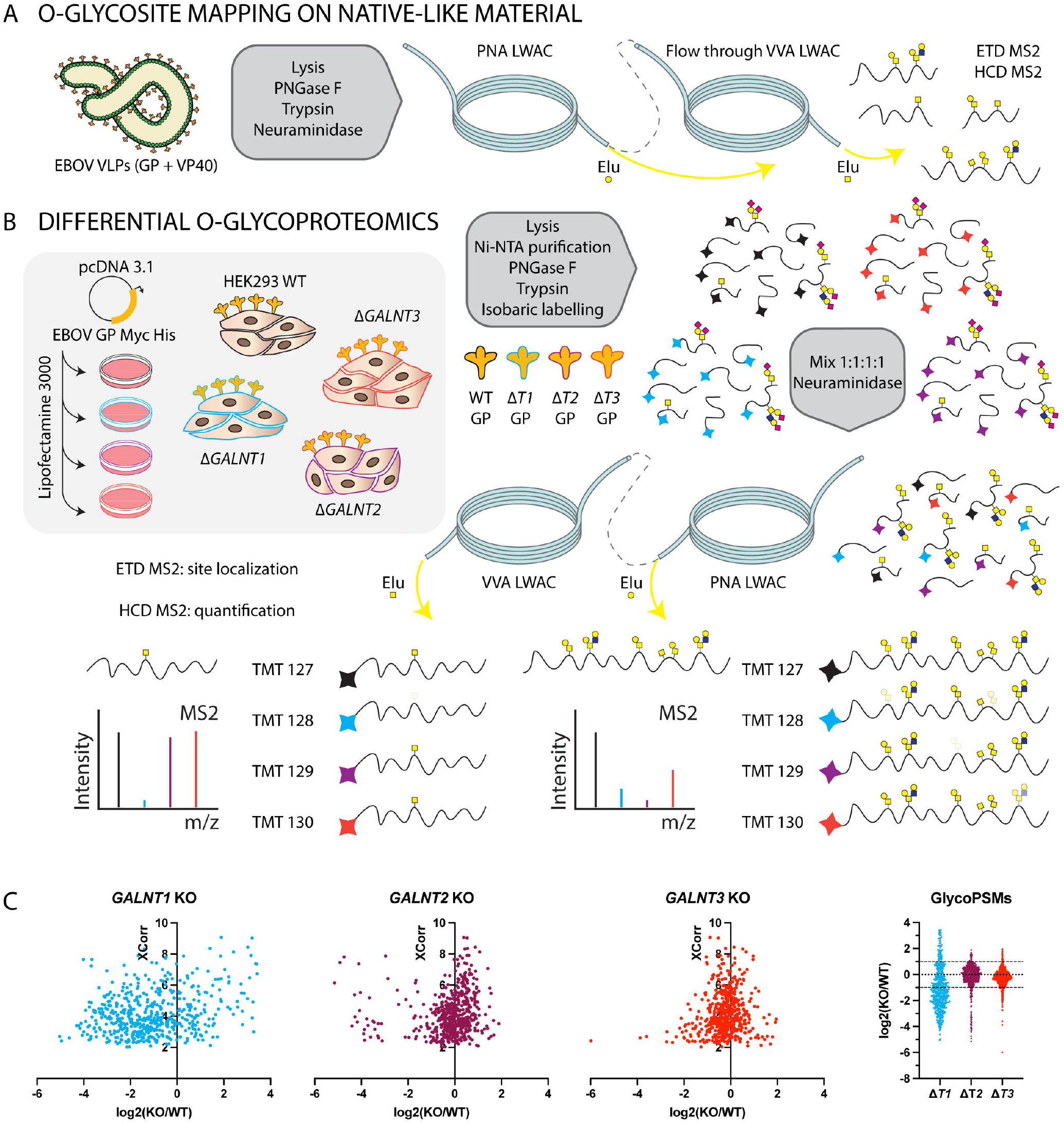
A strategy for Ebola virus O-glycoproteomics. (A) Approach for mapping O-glycosites on Ebola virus-like particles comprised of GP and VP40 proteins. (B) Approach for mapping differentially glycosylated O-glycosites on full-length recombinant GP expressed in *GALNT* KO cell lines. (C) The dot plots depict distribution of TMT quantification ratios compared to wild type (log_2_) for glycopeptide PSMs (peptide spectrum matches) identified in both PNA and VVA LWAC experiments plotted against XCorr (cross-correlation score) values.

**Figure 4.**
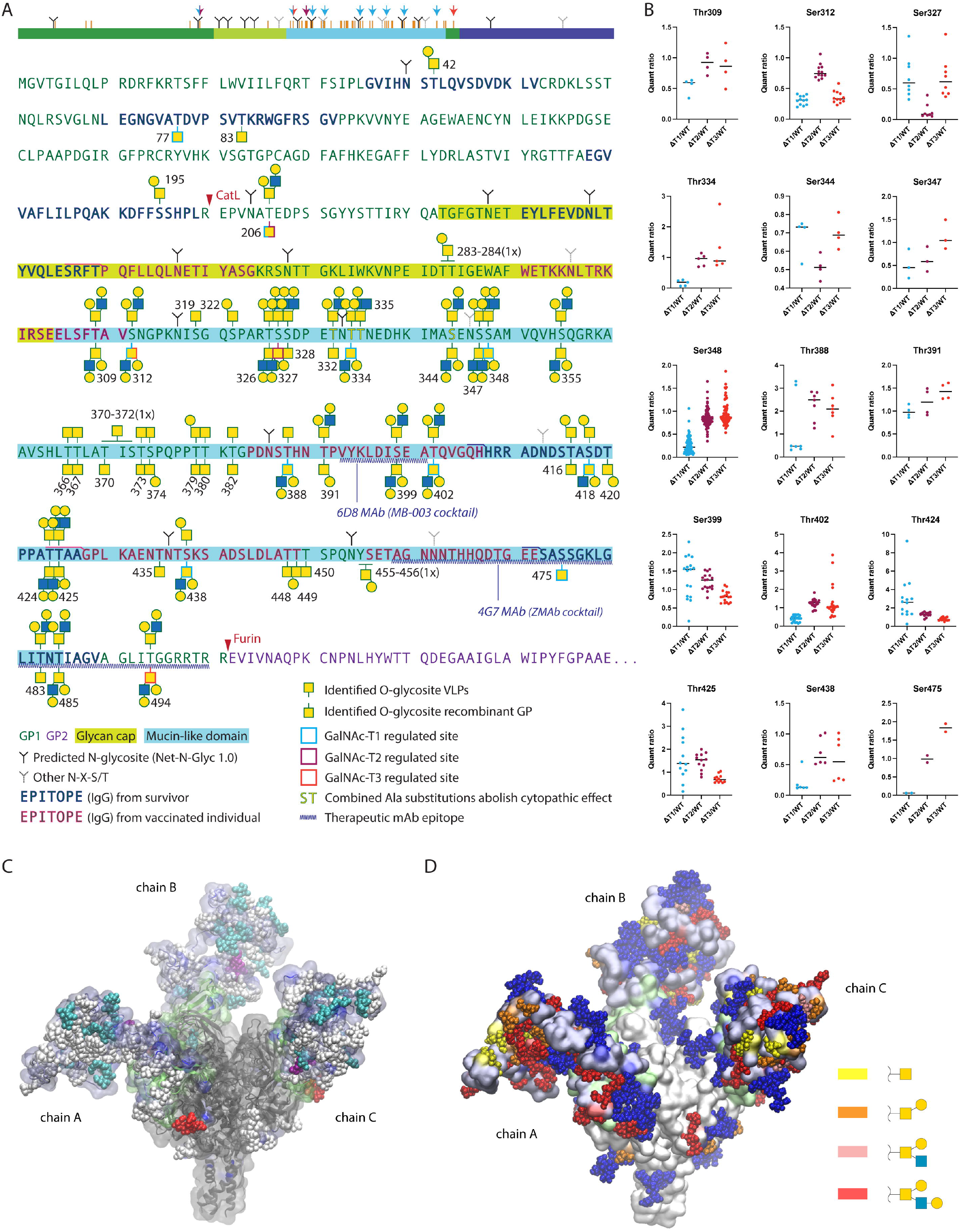
Mapping isoform-specific O-glycosites on EBOV GP. (A) Identified O-glycosites and most-complex unambiguously assigned site-specific structures are shown in the context of EBOV GP primary sequence, where VLP-derived sites are shown above the sequence, and recombinant GP-derived sites - below the sequence. Glycosites with median quantification ratios of singly glycosylated peptides below 0.5 were considered as isoform regulated and are outlined in cyan, maroon, and red for GalNAc-T1, GalNAc-T2, and GalNAc-T3, respectively. A simplified cartoon above summarizes the data, where orange bars represent all identified O-glycosites. Color coded arrows indicate isoform-regulated glycosites. Epitopes of protective antibodies derived from convalescent or vaccinated individuals, as well as epitopes for several therapeutic mAbs are annotated in the sequence (Heidepriem *et al*., 2020; Ponomarenko *et al*., 2014). Amino acids for which combined substitutions to Ala abolish cytopathic effect are also highlighted (Simon and Linstedt, 2018). (B) TMT quantification ratios of single site peptides, where each dot represents a separate PSM, and horizontal bars indicate median values. (C) Molecular modelling of EBOV GP with the identified O-glycans attached. The MLD was built *de novo* based on available cryo-EM/ET density maps (Beniac and Booth, 2017; Tran *et al*., 2014) of virion- and VLP-derived GP and combined with an atomic resolution structure of the GP lacking the MLD (PDB: 6HS4). The MLD is shaded in iceblue and the GCD is shaded in lime. GalNAc-T1, -T2, and -T3 regulated O-glycosites are highlighted in cyan, purple, and red, respectively. The remaining O-glycans are shown in white. Chain A contains VLP-derived glycosites, chain B – recombinant GP-derived glycosites, and chain C – combined maximum capacity. (D) Identified O-glycans are colored based on the longest site-specific structure identified, as indicated in the legend. Putative N-glycans were included in the model and are shown in blue.

Glycosites and site-specific structures identified on recombinant EBOV GP was in good agreement with those identified on VLPs, representing mature glycoproteins (Fig. 4A, Table 1). While many of the glycosites were identified on multi-glycosylated peptides, some of those sites were also covered by peptides only carrying a single O-glycan, possibly informing on the order of glycan addition. By quantifying the relative abundances of singly glycosylated peptides, we could evaluate the individual contributions of GalNAc-T1, -T2, and -T3 to specific sites (Fig. 3B, 3C, 4B). Non-redundant regulated glycosites could be identified for each isoform (Fig. 4A, 4B, Table 1) demonstrating that all three GalNAc-Ts contributed to complete glycosylation of EBOV GP. In addition, there were a few glycosites with contribution from several GalNAc-Ts, and we could also identify non-regulated sites, suggesting either preference by a different isoform, or non-selective glycosylation (Fig. 4B). GalNAc-T1 had by far the largest contribution, with selective regulation of eight O-glycosites, whereas GalNAc-T2 and GalNAc-T3 were responsible for one glycosite each (Fig. 3C, 4A). Moreover, knock out of GalNAc-T1 had a global effect on EBOV GP glycosylation and resulted in considerable changes of both downregulated and upregulated glycopeptides (Fig. 3C), suggesting GalNAc-T1 is a major regulator of EBOV GP glycosylation, supported by a notable molecular mass shift of the protein (Fig. S2). Interestingly, a big proportion of GalNAc-T1 regulated sites resided in close vicinity to N-glycosites, and most were found on de-amidated peptides, which likely were N-glycosylated before PNGase F treatment. Importantly, these glycosites were unambiguously localized to respective serines and threonines and often harbored complex core 1 and core 2 structures, excluding the possibility of artefactual GlcNAc “stumps” at N-glycosites (Table 1, Dataset S1, Fig. S3).

To visualize the identified O-glycosites in the context of mucin-like domain structure we used cryo-EM/ET density maps of virion- (11 Å) and VLP-derived GP (Beniac and Booth, 2017; Tran et al., 2014), as well as an atomic resolution structure of recombinant EBOV GP lacking the MLD (PDB: 6HS4, 2.05 Å), as a starting point to build the mucin-like domain *de novo*, while accommodating for the identified site-specific O-glycans (Fig. 4C, 4D). In addition, we performed a molecular dynamics simulation (100 ns) to model the behavior of the MLD, where the chain A of the trimer represents O-glycosylation of VLPs, chain B – that of recombinant GP, and chain C – maximum O-glycosylation capacity (Video S1-S4). We hereby present a putative model of how O-glycans could help shape the structure of this elusive domain (Fig. 4C, 4D).

### O-glycosylation of host cells modulates entry of pseudotyped EBOV

To investigate the effect of O-glycosylation on EBOV GP function, we performed two sets of experiments with EBOV GP pseudotyped recombinant vesicular stomatitis virus (VSV), lacking the gene encoding the G glycoprotein (VSVΔG) and only able to perform a single-replication cycle. To address GP-mediated entry to cells lacking O-glycan elongation or isoform-specific initiation, we infected the panel of glycoengineered cells with the wild type EBOV GP-VSV and quantified the luciferase reporter expression to evaluate the efficiency of viral infection. The assay revealed diminished entry into all mutant cell lines, suggesting host cell O-glycosylation may contribute to delayed propagation and decreased infectivity observed with live EBOV (Fig. 5A).

**Figure 5.**
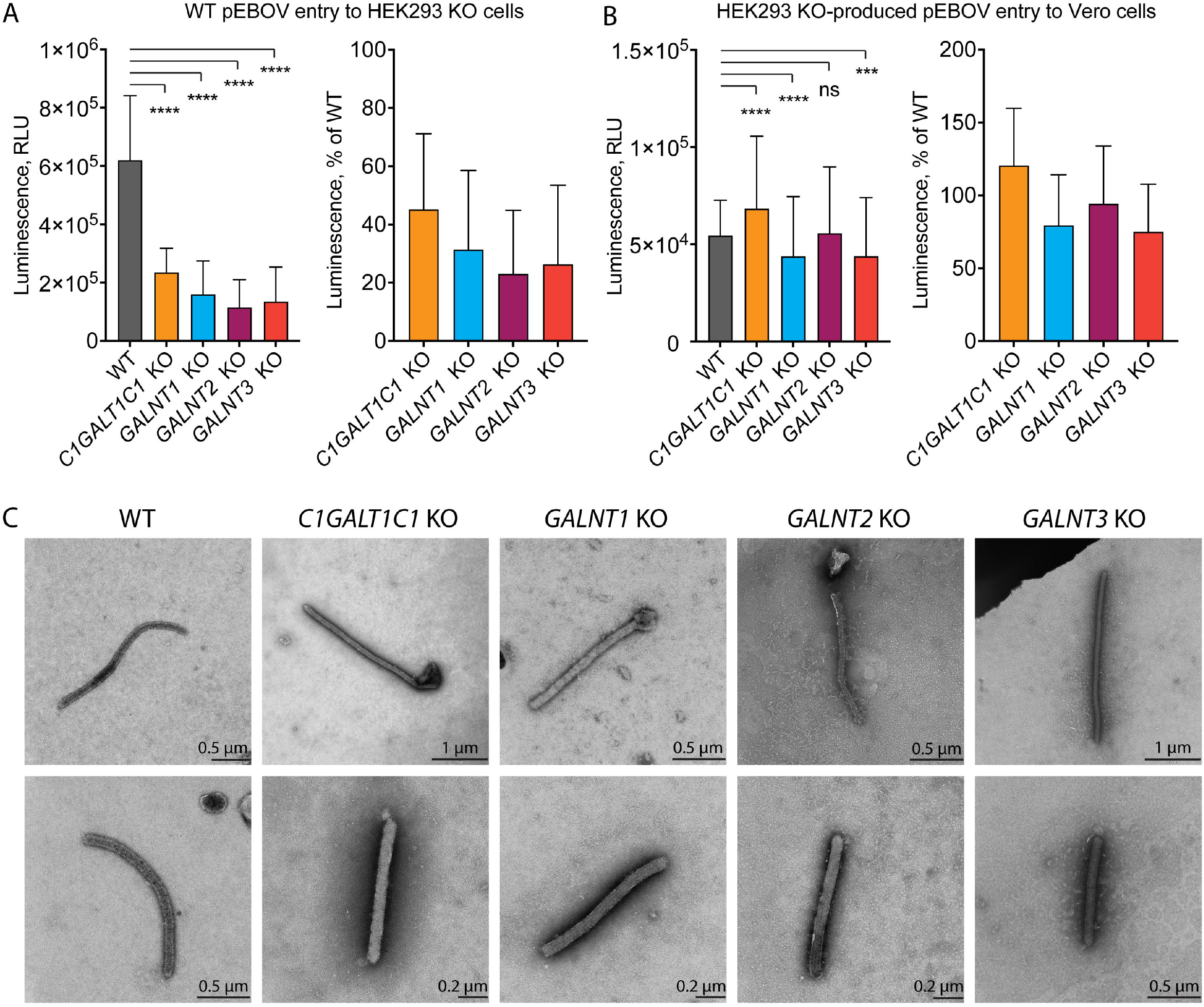
Influence of GalNAc-Ts on EBOV biology. (A) Entry of pEBOV-VSVΔGLuc to different HEK293 KO cells shown as reporter gene expression 24 hours post-infection. The data is shown as mean +SD of 5 biological replicates from 3 independent experiments. Two-way ANOVA followed by Dunnett’s multiple comparison test was used to evaluate differences from wild type (* *p* < 0.05, ** *p* < 0.01, *** *p* < 0.001, **** *p* < 0.0001). (B) Titers of HEK293 KO cell line-produced pEBOV-VSVΔGLuc harvested 14 hours post-infection and assayed on Vero cells by measuring reporter gene expression 24 hours post-infection. The data is shown as mean +SD of 8 biological replicates from 4 independent experiments. Two-way ANOVA followed by Dunnett’s multiple comparison test was used to evaluate differences from wild type (* *p* < 0.05, ** *p* < 0.01, *** *p* < 0.001, **** *p* < 0.0001). (C) TEM micrographs of Ebola virions produced in the panel of glycoengineered cell lines 24 hours post-infection. Negative staining with phosphotungstic acid was used for contrasting the specimens. Scale bar is indicated on each image.

To address the functionality of pseudovirus produced in the different knock out cell lines, the cells were transfected with EBOV GP encoding plasmid followed by infection with VSVΔG and titration of generated stocks on Vero cells. Pseudoviruses generated in the mutant cell lines exhibited similar titers to those in the wild type, though slightly lower titers were observed for *GALNT1* KO- and *GALNT3* KO-generated pEBOV, whereas the titer was slightly higher for *C1GALT1C1* KO-produced pEBOV (Fig. 5B). This suggests EBOV GP with altered O-glycosylation patterns is functional when presented on the VSV surface. To investigate whether particle morphology of infectious EBOV is affected by the O-glycosylation patterns, we performed transmission electron microscopy of virions present in cell culture supernatants of infected HEK293 cells 24 hours post-infection. We could detect characteristic rod-shaped virions in supernatants of all HEK293 KO cell lines (Fig. 5C). Aside from predominantly shorter *GALNT1* KO virions, we could not detect any gross changes in particle morphology. This does not exclude potential issues with particle functionality and ultrastructural studies would be needed to investigate GP distribution.

## Discussion

Glycosylation of viral glycoproteins plays diverse important roles in infectivity and immune shielding (Bagdonaite and Wandall, 2018). Studies on glycosylation of viral envelope proteins of highly pathogenic viruses are challenging due to several limitations, including difficulties and biosafety concerns in using the fully infectious viruses as test subjects. Furthermore, due to lack of technology to eliminate specific glycosylation sites without introducing large deletions or single amino acid mutations, and thereby changing the conformation of proteins, there has been a gap of knowledge regarding the effect of site-specific O-glycosylation. Here, we used a panel of genetically engineered cells in the context of natural EBOV infection to evaluate the consequences of global changes in O-glycan structures or downregulation of O-glycans at specific sites by eliminating isoforms of enzymes responsible for O-glycan initiation. In addition, we mapped the individual O-glycosites on EBOV GP and identified those GalNAc-T regulated subsets by differential O-glycoproteomics, with GalNAc-T1 having the most pronounced role. In addition, we used infection with native Ebola virus and observed a striking reduction in viral titers, not only for *C1GALT1C1* KO and *GALNT1* KO, but also for *GALNT2* KO and *GALNT3* KO cell lines. This would suggest involvement of specific O-glycosylation sites in EBOV pathogenesis, which has been suggested by large deletions of MLD, and by effects on recombinant EBOV GP-induced cell rounding by O-glycan truncation and loss of GalNAc-T1 (Simon and Linstedt, 2018). In addition, we have previously shown requirement of elongated O-linked glycans for efficient propagation of HSV-1, another heavily O-glycosylated virus (Bagdonaite et al., 2015). Experiments with EBOV GP pseudotyped VSV particles suggested contribution of all three investigated GalNAc-Ts, as well as O-glycan elongation of the host cells, to entry of wild type pseudovirus. This could both be related to O-glycosylation of several C-type lectins and TIM-1, as well as overall changes in the glycocalyx organization due to multiple affected substrates not directly involved in pathogenesis. For instance, glycan-glycan interactions have been suggested important for early HIV-1 interactions with the host cell (Spillings et al., 2022). It is important to note that epithelial cells will present a limited repertoire of C-type lectins, and there might be other not yet defined proteins mediating early interactions. While O-glycosylation defects did not affect GP incorporation into pseudotyped virions or their entry via clathrin-mediated endocytosis, it does not exclude detrimental effects on live virions, exhibiting entirely different morphology and entering via macropinocytosis, and is yet to be addressed. Furthermore, investigating the effect of O-glycosylation in cell types serving as primary entry points for EBOV infection *in vivo* would be advantageous.

While it is well known EBOV GP is heavily O-glycosylated, specific glycosites or the extent of glycan density within the MLD have not been described in detail. To address this question, we performed O-glycoproteomic analysis of EBOV VLPs to estimate the total capacity of O-glycosylation in cells expressing a complete set of GalNAc transferases, and identified 38-specific O-glycosites primarily within the glycan cap and the MLD, and could confirm the majority of those sites on recombinantly expressed GP with predominant core 1 and core 2 structures. By performing quantitative differential O-glycoproteomic analysis using *GALNT* KO cell lines, we found that GalNAc-T1 deficiency downregulated a large proportion of rather uniformly spaced O-glycosites and is expected to affect overall glycan density on the MLD, possibly also influencing its physical and immunological properties. GalNAc-T1 has previously been suggested to be responsible for MLD-induced cell rounding, and compound deletion of five specific Ser/Thr amino acids has achieved the same effect (Simon and Linstedt, 2018). Four of those amino acids are homologous in our investigated strain, and we found three of them O-glycosylated (Fig. 4A). Of those, Thr334 was selectively glycosylated by GalNAc-T1, and may be responsible for the MLD-induced cytopathic effect via a yet undefined mechanism.

Regulation of O-glycan initiation by competing GalNAc-Ts is not completely understood, and while there is no consensus sequence for O-glycosylation in general, or the individual isoforms, there are slight differences in amino acid context preference by the individual catalytic domains (Bagdonaite et al., 2021; de Las Rivas et al., 2019). Furthermore, most GalNAc-Ts are capable of long-range follow-up due to positioning by the lectin domain recognizing distant already glycosylated sites, where varying linker length between the two domains determines the range (de Las Rivas et al., 2017; Fritz et al., 2004; Pedersen et al., 2011). The N-glycosite proximal GalNAc-T1 regulated sites may be a result of the lectin domain-positioned O-glycan initiation, given GalNAc-T1 prefers to follow up 8-10 residues N-terminally from existing O-glycosites (Brokx et al., 2003; Tetaert et al., 2001), which would fit well with e.g., regulated Thr334 and Ser348 each spaced by 10 amino acids from respective Ser344 and Thr448.

A few O-glycosites have been identified close to cathepsin L cleavage site, with Thr206 regulated by GalNAc-T1 and GalNAc-T2, absence of which may affect processing in the endocytic compartment. Similarly, a GalNAc-T3 regulated site (Thr494) was found close to furin cleavage site, and may play a role in protein maturation. A number of immunodominant epitopes have been identified within the MLD and the glycan cap in animal studies, vaccine clinical trials and human survivors (Khurana et al., 2020; Sanchez-Lockhart et al., 2018; Wilson et al., 2000). Interestingly, rather consistent antibody signatures have been identified in the less glycosylated GP2, whereas there is little correlation of antibody epitopes in GP1 containing mucin-like domain and glycan cap between vaccinated and convalescent individuals (Heidepriem et al., 2020) (Fig. 4A). In non-human primates, vaccination approaches have generated the most immunodominant epitopes in the MLD, where class switched antibodies were also mapped (Sanchez-Lockhart *et al*., 2018). This may suggest variable antibody signatures to immunogens bearing differential O-glycosylation patterns due to distinct GalNAc-transferase repertoires in production cell lines and natural hosts (Bagdonaite *et al*., 2018). We found four O-glycosites (Ser475, Thr483, Thr485, and Thr494), two of them regulated, in the epitope region of 4G7 neutralizing antibody that is part of the ZMAb/ZMApp therapeutic antibody cocktails (Ponomarenko et al., 2014; Wilson *et al*., 2000) (Fig. 4A). Of those, Thr483 and Thr485 comprise epitopes for two other antibodies. In addition, Ser399, which we find glycosylated, is part of the epitope for 6D8 monoclonal antibody, which is included in MB-003 cocktail (Ponomarenko *et al*., 2014) (Fig. 4A). Understanding these signatures in relation to GalNAc-T repertoires and glycosite occupancy would be useful for understanding principles of neutralization and could help generate desirable immune responses by glycoengineering the immunogens.

## Acknowledgements

This research was funded by the European Commission (GlycoSkin H2020-ERC), Lundbeck Foundation (R219-2016-545; R313-2019-869), Danish National Research Foundation (DNRF107), Neye Foundation, Novo Nordisk Foundation, and University of Padua (DOR 2021). This study has also received funding from the Innovative Medicines Initiative 2 Joint Undertaking (JU) under grant agreement No. 823666 and by the EU/CEPHEID Innovative Medicines Initiative Joint Undertaking VHFMoDRAD grant No. 823666. The JU receives support from the European Union’s Horizon 2020 research and innovation programme and EFPIA and CEPHEID EUROPE SAS. We acknowledge the Core Facility for Integrated Microscopy, Faculty of Health and Medical Sciences, University of Copenhagen.

## Author contributions

Conceptualization, I.B., C.S., A.M., and H.H.W.; Methodology, I.B., S.A., M.M., M.F., C.S., and S.Y.V.; Software, M.F. and S.Y.V.; Validation, I.B.; Formal Analysis, I.B., S.A., M.M., M.F., and S.Y.V.; Investigation, I.B., S.A., M.M., M.F., S.Y.V., C.S., A.M. and H.H.W.; Resources, M.F., Y.N., S.Y.V., C.S., A.M., and H.H.W.; Data Curation, I.B. and S.Y.V.; Writing – Original Draft Preparation, I.B. and H.H.W.; Writing – Review & Editing, I.B., S.A., M.M., M.F., Y.N., S.Y.V., C.S., A.M., and H.H.W.; Visualization, I.B., M.F., and H.H.W.; Supervision, C.S., A.M., and H.H.W.; Project Administration, I.B. and H.H.W.; Funding Acquisition, I.B., C.S., A.M., and H.H.W. All authors have read and agreed to the published version of the manuscript.

## Materials and methods

### Cells, plasmids and viruses

The cell lines used were HEK293T/17 (HEK293T, ATCC^®^ CRL-11268™), HEK293 WT (Sigma Cat. Nr. 85120602), HEK293 *GALNT1* KO (Narimatsu et al., 2019b), HEK293 *GALNT2* KO (Narimatsu *et al*., 2019b), HEK293 *GALNT3* KO (Narimatsu *et al*., 2019b), *C1GALT1C1* KO (Steentoft et al., 2013) bearing truncated O-glycans (SC), VERO (VERO-ccl81, ATCC^®^ CCL-81™), and Vero-E6 cells (ATCC, CRL-1586™). All cell line were maintained in Dulbecco’s Modified Eagle’s Medium (Life Technologies Cat. Nr. 419660), supplemented with 10 % v/v of heat-inactivated Fetal Bovine Serum (FBS, Life Technologies Cat. Nr. 10500) and incubated at 37° C with 95 % humidity and 5% CO_2_.

Zaire ebolavirus (ZEBOV isolate Ebola virus/H.sapiens-wt/SLE/2014/Makona-G3838) was propagated in Vero-E6 cells in T75 culture flasks. At day six post-infection, virus particles in cell culture supernatants were harvested and clarified from cell debris by centrifugation at 1,500 rpm for 10 min. The virus was then aliquoted in Eppendorf tubes, titrated in Vero-E6 cells cultured in 16-well chambered slides (Thermo Fisher Scientific) as described below and stored at -80 °C. The plasmid expressing the Zaire ebolavirus (ZEBOV) Glycoprotein (isolate Ebola virus/H.sapiens-wt/SLE/2014/Makona-G3838), pcDNA3.1-ZEBOV-GP, was kindly provided by Stefan Pöhlmann (Deutsches Primatenzentrum GmbH, Germany). To generate the *myc*- and His-tagged version of the protein, GP sequence was subcloned to a pcDNA™3.1/*myc*-His(-) plasmid. The plasmid expressing Zaire ebolavirus matrix protein VP40 fused in-frame with the green fluorescent protein (GFP) (pVP40-GFP) was a gift from Christopher Basler (Mount Sinai School of Medicine, New York, USA). The plasmid expressing the Vesicular Stomatitis Virus (VSV) glycoprotein (pVSV-G) was previously described (Salata et al., 2009). The recombinant VSV encoding the luciferase in place of the VSV-G gene (VSVΔG-Luc) was provided by Michael Whitt, University of Tennessee, USA.

### EBOV infection

HEK293 wild type and KO cells (WT, *C1GALT1C1* KO, *GALNT1* KO, *GALNT2* KO, and *GALNT3* KO) were seeded in T25 culture flasks in DMEM containing 10 % FBS and 50 U/mL penicillin/streptomycin (Thermo Fisher Scientific Cat. Nr. 15070). Twenty-four hours post-seeding, cells were infected with ZEBOV at an MOI of 0.1 in infection medium (DMEM containing 2 % FBS). One hour post-infection, input virus were discarded and cells were washed once with infection medium, added fresh complete medium and incubated further at 37 °C, 5 % CO_2_. At one and six days post-infection, virus particles in the culture supernatant (progeny viruses) were harvested and clarified from cell debris by centrifugation at 1,500 rpm for 10 min. Titration of progeny viruses from all HEK293 cells was done in Vero-E6 cells cultured in 16-well chambered slides (Thermo Fisher Scientific), as described below. For RNA extraction, cells and cell culture supernatants were harvested in TRIzol reagent (Thermo Fisher Scientific) at day one and six post-infection. For immunofluorescence experiments, HEK293 wild type and knockout cells were grown on 12-well chambered glass slides and infected with ZEBOV at an MOI of 0.01 in DMEM containing 2 % FBS for 1 h at 37°C, 5 % CO_2_. Thereafter input virus was discarded and new complete medium was added to each well and incubated further at 37 °C, 5 % CO_2_. At 24 hours post-infection the slides were inactivated in acetone twice, first for 10 min and then transferring to another jar with new acetone and incubating for 30 min at room temperature.

### Fluorescent focus forming assay

Infectivity of progeny viruses produced from HEK293 wild type and knockout cells were determined using infectious focus forming assay. Briefly, Vero-E6 cells cultured in 16-well chambered slides were infected with ten-fold serially diluted virus for 1 h at 37 °C, 5 % CO_2_. Input virus were then discarded and fresh complete medium was added to the cells. At 24 hours post-infection, cells were fixed in acetone for 20 min. Cells were permeabilized in PBS that contained 0.1 % Triton X-100 for 5 min. Slides were then incubated with rabbit anti-EBOV GP primary antibody (produced by Agrisera upon request) that was diluted in 150 mM NaCl for 1 h, washed three times in PBS and further incubated with FITC-conjugated goat anti-rabbit IgG (Thermo Fisher Scientific) as a secondary antibody for 30 min. All incubations were carried out at 37 °C. DAPI (4’,6-diamidino-2-phenylindole dihydrochloride) was used to stain cell nuclei.

### RT-qPCR

Samples from infected cells and cell culture supernatants were extracted using TRIzol reagent (Thermo Fisher Scientific). At day one and six post-infection, cell culture supernatants were harvested by mixing with TRIzol LS reagent (Thermo Fisher Scientific) at a ratio of 1 to 3. Cells in T25 culture flask were harvested in 1 mL TRIzol reagent. Prior to RNA extraction, TRIzol inactivated cells and cell culture supernatants were subjected to chloroform treatment, followed by RNA extraction using PSS magLEAD 12gC machine (Precision System Science Co.) and eluted the RNA with 100 µL elution buffer. Purified samples were stored at -80°C, pending analysis.

The ZEBOV RT-PCR assay was performed in a 25 µL reaction mixture containing TaqMan Fast Virus 1-step master mix (Thermo Fisher Scientific), 5 µl template RNA, 900 µM of each primer and 200 µM of TaqMan probe. The forward primer 5’-ATGGGCTGAAAAYTGCTACAATC and reverse primer 5’-CTTTGTGMACATASCGGCAC as well the probe FAM-CTACCAGCAGCGCCAGACGGGA-TAMRA were for amplification of ZEBOV GP gene. Amplification and detection of the vRNA was performed in a StepOne Plus real-time PCR machine (Applied Biosystems). The cycling conditions were as following: 50 °C for 5 min; 45 cycles of 95 °C for 20 sec and 95 °C for 3 sec and 60 °C for 30 sec. Human beta-actin was used as an endogenous gene control to confirm the integrity of the extraction reagent and RNA recovery as well as to normalize the levels of intracellular viral RNAs.

### Ebola virus-like particle production

Zaire ebolavirus-Like particles (EBOVLPs) were produced by transfection of HEK293T cells with a plasmid expressing the EBOV matrix protein VP40 fused to GFP (VP40-GFP) along with a construct expressing either the ZEBOV GP – strain Makona, or the VSV envelope G glycoprotein. Cells seeded in 100 mm plates were transfected with 11.5 µg of pVP40-GFP and along with 0.5 µg of pcDNA3.1-ZEBOV-GP or pVSV-G by calcium-phosphate (Calistri et al., 2009). Cell culture supernatants were collected 48 hours after transfection, cell debris were cleared by centrifugation (1,200 rpm for 7 minutes at 4 °C). EBOVLPs production was confirmed by cytofluorimetric analyses of transduced Vero CCL81 cells as previously described (Salata et al., 2018). Next, VLPs were purified and concentrated by two ultracentrifugation steps at 27,000 rpm for 2 hours in 20 % sucrose cushion, then aliquoted and stored at -80 °C.

### Pseudotyped virus production and titration

ZEBOV GP pseudotyped VSVΔG-Luc (GP-pseudotyped virus) was generated as previously described (Salata et al., 2015). Briefly, HEK293T cells were transfected by calcium-phosphate protocol with 16 μg of pcDNA3.1-ZEBOV-GP plasmid and, 24 h later, infected with the recombinant VSVΔG-Luc virus at the multiplicity of infection (MOI) of 4 fluorescent focus-forming units (FFU)/cell. After 24 h, the pseudotyped virus was harvested and stored at −80°C. Viral titer was evaluated as FFU/mL using the anti-VSV-Matrix protein 23H12 antibody (Kerafast Inc.) to detect infected cells.

### EBOV-Pseudotype (pEBOV_KO) Production on HEK293 knockout cell lines

HEK293 were seeded in poly-Lysine-coated 6-well plates and transfected with the pcDNA3.1-ZEBOV-GP plasmid as above described. Six h later, cells were washed and fresh medium added. The day after, cells were infected for 1 h with the VSVΔG-Luc virus at MOI 4 FFU/cell, washed twice in PBS and incubated in complete medium for 14 h. Next, medium was recovered and centrifuged at 3,500 rpm at 4° C for 6 minutes to remove cellular debris. Then, aliquots were made and stored at -80° C for further use. The viral progeny production was evaluated infecting semi-confluent Vero-CCL-81 cells seeded in 96 well plates. After 1 h cells were washed in PBS and cultured for additional 24 h before the evaluation of the Luc expression using the Steady-Glo^®^ Luciferase reagent (Promega), which were added to the cell for 15 minutes. The luminescence was detected with a VICTOR Multilabel Plate Reader (PerkinElmer) as relative-luminescence unit (RLU, normalized on a scale factor of 1 second).

### pEBOV infection on HEK293 knockout lines

One day before the assay, 2×10^4^ HEK293 knockout cells were seeded in poly-Lysine-coated 96-well plates. The day after, cells were washed with PBS and infected with pEBOV at MOI 0.01 for 1 hour. Then, cells were washed with PBS and new complete medium was added. After 24 h the reporter gene expression was detected, as above described, to evaluate the efficiency of infection.

### Transfection

For immunofluorescence experiments the panel of HEK293 cells grown on poly-Lysine (Sigma-Aldrich Cat. Nr. P6282) pre-coated (2.5 μg/mL in MQ H_2_O 30 min at room temperature (RT)) glass cover slips in 24-wells were transfected with 0.5 μg/well of pcDNA™3.1/*myc*-His(-) plasmid encoding *myc*- and His-tagged full-length EBOV GP (Zaire ebolavirus isolate Ebola virus/H.sapiens-wt/SLE/2014/Makona-G3838) using Lipofectamine™ 3000 (Thermo Fisher Scientific Cat. Nr. L3000) according to manufacturer’s instructions. Non-transfected cells were used as controls. At 48 hours post-transfection the cells were washed 3x with Hanks⍰ Balanced Salt solution (HBSS, Sigma-Aldrich Cat. Nr. H8264) and fixed with 4 % PFA in PBS (Ampliqon Cat. Nr. AMPQ44154.1000) for 10 min at RT followed by 3x more washes. If needed, the cells were permeabilized with 0.3 % Triton X-100 in HBSS for 3 min followed by 3x washes with PBS and blocked with 2.5 % BSA in PBS, 0.03 % sodium azide at 4 °C until staining. For Western blotting experiments the panel of HEK293 cells grown in 6-wells were transfected with 2 μg/well of the plasmid and harvested at 48 hours post-transfection. The cells were washed with PBS and lysed in modified RIPA buffer (50 mM Tris pH 7.5, 150 mM NaCl, 1 % NP-40, 0.1 % Na deoxycholate, 1 mM EDTA) supplemented with protease inhibitor cocktail (cOmplete™, EDTA-free, Roche Cat. Nr. 11873580001) and phosphatase inhibitors for 30 min at 4 °C with agitation, followed by sonication using a sonic probe on ice. The lysates were centrifuged at 10000 x g for 10 min at 4 °C and the supernatants used for subsequent Western blotting. For protein purification HEK293 WT, *GALNT1* KO, *GALNT2* KO, and *GALNT3* KO cells were grown in 3x P150 dishes each and transfected with 23 μg/dish of the plasmid and harvested at 48 hours post-transfection. The plates were chilled on ice, washed with ice-cold PBS twice and cells gently scraped in ice-cold PBS followed by centrifugation at 500 x g for 10 min at 4 °C. After decanting most of PBS, pellets were dislodged and pooled for each cell type, followed by centrifugation at 800 x g for 10 min at 4 °C. PBS was removed, and pellets stored at -80 °C.

### EBOV GP purification

Cell pellets were thawed on ice in 0.1 % RapiGest (Waters Cat. Nr. 186001861) in Equilibration buffer (25 mM Tris-HCl 300 mM NaCl pH 7.5) supplemented with protease inhibitor cocktail (cOmplete™, EDTA-free, Roche Cat. Nr. 11873580001) followed by sonication using a sonic probe on ice. His-tagged proteins were purified using HisPur™ Ni-NTA resin (Thermo Fisher Scientific Cat. Nr. 88222) according to manufacturer’s instructions (batch protocol). Briefly, the lysates were cleared by centrifugation at 10000 x g for 10 min at 4 °C and applied to pre-equillibrated HisPur™ Ni-NTA resin followed by a 1-hour incubation on an end-over-end rotator. The resin was washed 3x 5 min with Wash buffer (25 mM Tris-HCl 300 mM NaCl pH 7.5, 10 mM imidazole), and bound proteins eluted in Elution buffer (25 mM Tris-HCl 300 mM NaCl pH 7.5, 250 mM imidazole). Protein concentration was measured using Pierce™ 660 nm Protein Assay Kit (Thermo Fisher Scientific Cat. Nr. 22662) and 400 μg of each cell type-derived protein taken for subsequent MS sample preparation.

### Immunofluorescence

Fixed cover slips or teflon-coated slides were incubated with primary antibodies diluted in 2.5 % BSA (Sigma-Aldrich Cat. Nr. A3294) in PBS, 0.03 % sodium azide over night at 4 °C: rabbit anti-EBOV GP (1:300, produced by Agrisera upon request), goat anti-E-cadherin (1:200, R&D Systems Cat. Nr. AF648), rabbit anti-Myc (1:200 Abcam Cat. Nr. ab152146) or undiluted hybridoma supernatants (mouse anti-GalNAc-T1 (4D8), mouse anti-GalNAc-T2 (4C4), mouse anti-GalNAc-T3 (2D10)), followed by 3x washes with PBS and 1-hour incubation at RT with secondary antibodies: donkey anti-rabbit IgG AF546 (1:500), donkey anti-goat IgG AF488 (1:500), goat anti-rabbit IgG AF488 (1:500), goat anti-mouse IgG AF594 (1:500), all from Thermo Fisher Scientific. After 3x washes with PBS cover slips were incubated with 0.1 μg/mL DAPI solution for 4 min followed by 3x washing and mounting with Prolong Gold antifade-reagent (Thermo Fisher Scientific Cat. Nr. P36930). Teflon-coated slides were mounted using Prolong Gold antifade-reagent with DAPI (Thermo Fisher Scientific Cat. Nr. P36935). Fluorescence micrographs and z-stacks were obtained on a Zeiss LSM710 confocal microscope. Images were assembled using Adobe Photoshop, Adobe Illustrator or Zeiss ZEN Lite software.

### Western Blotting

30 μg of protein extracts were divided into two aliquots, one of which was treated with 2 U of PNGase F (Roche Cat. Nr. 11365177001) at 37 °C for 1 hour. Samples were then mixed with 4x NuPAGE sample buffer (Thermo Fisher Scientific Cat. Nr. B007) and 10 mM DTT, heat denatured (95 °C 5 min), and separated on Novex 4-12 % gradient gel (Bis-Tris) (Thermo Fisher Scientific Cat. Nr. NP0329) in 1x NuPAGE MES running buffer (Thermo Fisher Scientific Cat. Nr. NP0002), followed by transfer onto nitrocellulose membrane in 20 % MeOH in running buffer at 320 mA for 1 hour. Membranes were blocked with 5 % skim milk in TBS-T and blotted with rabbit anti-EBOV GP (1:800, produced by Agrisera upon request) antibody over night at 4 °C, followed by goat anti-rabbit Igs-HRP (1:4000, DAKO Cat. Nr. P0448) for 1 hour at RT. Membranes were developed using Pierce™ ECL Kit (Thermo Scientific) and visualized using the ImageQuant LAS4000 system.

### Transmission electron microscopy

Cell culture supernatants from HEK293 WT and different KO cell lines harvested at 24 hours post-infection were inactivated by mixing 1:1 with 2.5 % freshly prepared glutaraldehyde and incubating for 1 h at room temperature. Samples were applied to copper grids (Gilder Grids, G400) and negative stained with phosphotungstic acid. Grids were analyzed using a CM100 transmission electron microscope (FEI/Philips) equipped with TWIN objective lens and a side-mounted Olympus Veleta camera with a resolution of 2048 × 2048 pixels (2K x 2K). Images were recorded using ITEM software.

### O-glycoproteomic sample preparation

For differential O-glycoproteomic analysis, 400 μg of each cell type-derived protein in equal volumes of Elution buffer were diluted to 1 mL using 0.5 % RapiGest in 250 mM AmBic (ammonium hydrocarbonate) and H_2_O resulting in a final concentration of 0.1 % RapiGest and 50 mM AmBic. Proteins were reduced by adding up to 5 mM dithiothreitol (DTT, Sigma-Aldrich Cat. Nr. D0632) and incubating at 60 °C for 45 min followed by alkylation with 10 mM iodoacetamide (IAA, Sigma-Aldrich Cat. Nr. I1149) at RT for 30 min in darkness. The proteins were then treated with 8 U PNGase F (Roche Cat. Nr. 11365177001) at 37 °C for 3 hours followed by 19 μg/sample of trypsin (Roche Cat. Nr. 11418025001) at 37 °C for 13 hours. Digests were acidified with trifluoracetic acid and peptides purified using Sep-Pak (1cc) C18 cartridges (Waters Cat. Nr. WAT023590) (1x CV MeOH; 1x CV 50 % MeOH 0.1 % FA; 3x CV 0.1 % TFA; load twice; 3x CV 0.1 % FA; elute in 2x CV of 50 % MeOH 0.1 % FA (CV = column volume)). Peptide concentrations were measured using NanoDrop A205. 200 μg of peptides from each sample was taken, most of solvent evaporated using SpeedVac then added up to 1 mL H_2_O and freeze dried. Dried peptides were reconstituted in 100 μL TEAB and labeled with TMTsixplex™ reagents (TMT 127, TMT 128, TMT 129, TMT 130, Thermo Fisher Scientific Cat. Nr. 90061) according to manufacturer’s instructions. 1 % of each reaction was mixed, dried and submitted to LC-MS analysis for a ratio check. After confirming equal ratios all four channels were mixed and dried with SpeedVac. The peptide mix was then reconstituted in 50 mM sodium acetate pH 5 and treated with 0.25 U/mL *Clostridium perfringens* neuraminidase (Sigma-Aldrich Cat. Nr. N3001) at 37 °C for 3 hours. For O-glycoproteomic analysis of EBOV VLPs 300 μL of VLPs were diluted with 0.1 % RapiGest in 50 mM AmBic up to 1.3 mL, then sonicated using a sonic probe on ice, heated at 80 °C for 10 min and protein concentration measured using Pierce BCA™ Assay Kit. Approximately 800 μg of VLP lysate was then reduced and alkylated as described above and treated with 5 U PNGase F (Roche Cat. Nr. 11365177001) at 37 °C for 12 hours followed by 12 hours with 11 μg of trypsin. PNGase F treatment was then repeated followed by a 2-hour incubation with 4 μg of trypsin. Peptides were purified as described above, concentrated by SpeedVac and treated with 0.15 U/mL neuraminidase at 37 °C for 3 hours.

### LWAC enrichment

Sequential PNA and VVA LWAC enrichment was performed as previously described (Bagdonaite *et al*., 2015). Elution fractions were desalted using self-made Stage Tips (C18 sorbent from Empore 3M) and submitted to LC-MS and HCD/ETD-MS/MS.

### Mass spectrometry analysis

LC MS/MS site-specific O-glycopeptide analysis of was performed on EASY-nLC1200 UHPLC (Thermo Fisher Scientific) interfaced via nanoSpray Flex ion source to an Orbitrap Fusion Lumos Tribrid MS (Thermo Fisher Scientific) or an on EASY-nLC1000 UHPLC (Thermo Fisher Scientific) interfaced via a PicoView nanoSpray ion source (New Objectives) to Orbitrap Fusion mass spectrometer (Thermo Fisher Scientific). The nLC was operated in a single analytical column set up using PicoFrit Emitters (New Objectives, 75 mm inner diameter) packed in-house with Reprosil-Pure-AQ C18 phase (Dr. Maisch, 1.9-mm particle size, 19-21 cm column length). Each sample was injected onto the column and eluted in gradients from 3 to 32 % B in 95 min, from 32 to 100 % B in 10 min and 100 % B in 15 min at 200 nL/min (Solvent A, 100 % H_2_O; Solvent B, 80 % acetonitrile; both containing 0.1 % (v/v) formic acid, EASY nLC-1200) and from 3 to 25 % B in 95 min, from 25 to 80 % B in 10 min and 80 % B in 15 min at 200 nL/min (Solvent A, 100 % H_2_O; Solvent B, 100 % acetonitrile; both containing 0.1 % (v/v) formic acid, EASY nLC-1000).

A precursor MS1 scan (m/z 350–1,700) was acquired in the Orbitrap at the nominal resolution setting of 120,000, followed by Orbitrap HCD-MS2 and ETD-MS2 at the nominal resolution setting of 50,000 of the five most abundant multiply charged precursors in the MS1 spectrum; a minimum MS1 signal threshold of 50,000 was used for triggering data-dependent fragmentation events. For EBOVLP samples, stepped collision energy +/-5 % at 27 % was used for HCD MS/MS fragmentation and charge dependent calibrated ETD reaction time was used with CID supplemental activation at 30 % collision energy for ETD MS/MS fragmentation. For TMT-labelled samples, stepped collision energy +/-5 % at 45 % was used for HCD MS/MS fragmentation and charge dependent calibrated ETD reaction time was used with CID supplemental activation at 30 % collision energy for ETD MS/MS fragmentation.

For the site-specific glycopeptide identification, the corresponding HCD MS/MS and ETD MS/MS data were analyzed by Proteome discoverer 1.4 Software (Thermo Fisher Scientific) for EBOVLP samples or Proteome discoverer 2.2 software (Thermo Fisher Scientific) for EBOV GP samples using Sequest HT as a searching engine. Carbamidomethylation at cysteine was used as fixed modification and oxidation at methionine, asparagine deamidation, and HexNAc, Hex1HexNAc1, Hex1HexNAc2, and Hex2HexNAc2 at serine/threonine/tyrosine were used as variable modifications. Precursor mass tolerance was set to 10 ppm and fragment ion mass tolerance was set to 0.02 Da. Data were searched against the human-specific UniProt KB/SwissProt-reviewed database downloaded on January, 2013 and construct-dependent viral protein sequence databases. All spectra of interest were manually inspected and validated to prove the correct peptide identification and glycosite localization. The mass spectrometry proteomics data have been deposited to the ProteomeXchange Consortium via the PRIDE (Perez-Riverol et al., 2019) partner repository with the dataset identifier PXD033994.

### Molecular Modeling

The initial three-dimensional model of the fully glycosylated Ebola virus GP was built using the graphical interface of YASARA (Krieger and Vriend, 2014). The model was based on the crystal structure of Ebola GP (PDB entry 6hs4, resolution 2.05 Å), which lacks most of the residues in the range 294-501. This amino acid sequence – which includes the MLD – was built *de novo* and the N-glycans and O-glycans (taken from an in-house 3D library) were attached to the protein based on information shown in Fig. 4. The glycopeptide was relaxed by MD simulation and then subsequently fitted into the crystal structure, taking into account partly resolved parts (residues 302-310 and 471-478) and guided by available cryo-EM/ET density maps of virion- and VLP-derived GP (Beniac and Booth, 2017; Tran *et al*., 2014). Other missing loops in PDB entry 6hs4 were also subsequently modeled and N- and O-glycans attached. Single amino acids mutations were introduced in order to match the target sequence (isolate Ebola virus/H.sapiens-wt/SLE/2014/Makona-G3838). The trimeric 3D structure was finally adjusted so that chain A contains VLP-derived glycosites, chain B – recombinant GP-derived glycosites, and chain C – combined maximum capacity. Since each chain contained up to 46 O-glycans and 17 N-glycosylation sites, it was instrumental to cross-check the glycosylation pattern in the molecular system during the building process and prior to simulation using Conformational Analysis Tools (CAT).

Finally, the trimeric glycoprotein was solvated in 0.9 % NaCl solution (0.15 M) and simulations were performed at 310 K using the AMBER14 force field (He et al., 2020; Kirschner et al., 2008; Maier et al., 2015). The box size (approx. 185Åx185Åx185Å, 643911 atoms) was rescaled dynamically to maintain a water density of 0.996 g/mL. Simulations were performed at a rate of 4 ns/day using YASARA with GPU acceleration in ‘fast mode’ (4 fs time step) (Krieger and Vriend, 2015) on ‘standard computing boxes’ equipped e.g. with one 12-core i9 CPU and NVIDIA GeForce GTX 1080 Ti.

Conformational Analysis Tools (CAT, http://www.md-simulations.de/CAT/) was used for analysis of trajectory data, general data processing and generation of scientific plots. VMD (Humphrey et al., 1996) was used to generate molecular graphics.

## Supplemental information titles and legends

**Dataset S1. Identified O-glycopeptides**. The different spreadsheets include EBOV GP-derived O-glycopeptides identified in recombinant GP and VLP samples upon sequential PNA and VVA LWAC enrichment. Glycosite assignments have been manually validated.

**Figure S1. GalNAc-T expression in HEK293 KO cell lines**. Cells transfected with plasmid encoding EBOV GP Myc His were fixed and permeabilized 48 hours post-transfection and co-stained for Myc-Tag (green) and GalNAc-T1 (A), GalNAc-T2 (B), and GalNAc-T3 (C) (red). Scale bar 10 μm.

**Figure S2. EBOV GP expression in HEK293 WT and KO cell lines**. HEK293 WT (WT), *C1GALT1C1* KO (SC), *GALNT1* KO (ΔT1), *GALNT2* KO (ΔT2), and *GALNT3* KO (ΔT3) cells were transfected will a plasmid encoding full length Myc- and His-tagged EBOV GP. 48 hours post-transfection the cells were lysed, and 15 μg of extracts treated with 2 U of PNGase F for 1 hour at 37 °C or incubated without treatment. Separated proteins were transferred onto a nitrocellulose membrane and probed for EBOV GP.

**Figure S3. Examples of O-glycosites localized close to/within N-glycan sequons carrying Hex2HexNAc2 modifications**. Structures are depicted as core 2 based on O-glycoprofiling of HEK293 cells, where elongated core 1 structures matching the same composition are not detectable (de Haan *et al*., 2022). Amino acids carrying modifications are highlighted in red, where identified O-glycosites are also in bold. TMT - tandem mass tag, oxid - methionine oxidation, deam - asparagine deamidation.

**Video S1. MD simulation of O-glycosylated EBOV GP with isoform-specific glycosites highlighted**. A section of a molecular dynamics (MD) trajectory (covering a timescale of about 60 ns) is shown. The MLD is shaded in iceblue and the GCD is shaded in lime. GalNAc-T1, -T2, and -T3 regulated O-glycosites are highlighted in cyan, purple, and red, respectively. The remaining O-glycans are shown in white. Chain A contains VLP-derived glycosites, chain B – recombinant GP-derived glycosites, and chain C – combined maximum capacity. Putative N-glycans are shown in blue.

**Video S2. 3D model of O-glycosylated EBOV GP with isoform-specific glycosites highlighted**. Last snapshot of the timescale covered in Video S1.

**Video S3. MD simulation of O-glycosylated EBOV GP with O-glycans colored by structure**. A section of a molecular dynamics (MD) trajectory (covering a timescale of about 60 ns) is shown. The MLD is shaded in iceblue and the GCD is shaded in lime. Identified O-glycans are colored based on the longest site-specific structure identified, as indicated in Fig. 4D legend. Putative N-glycans are shown in blue.

**Video S4. 3D model of O-glycosylated EBOV GP with O-glycans colored by structure**. Last snapshot of the timescale covered in Video S3.

