## Supplementary figures and images for "Isoform-specific O-glycosylation dictates Ebola virus infectivity"

### Figure S1

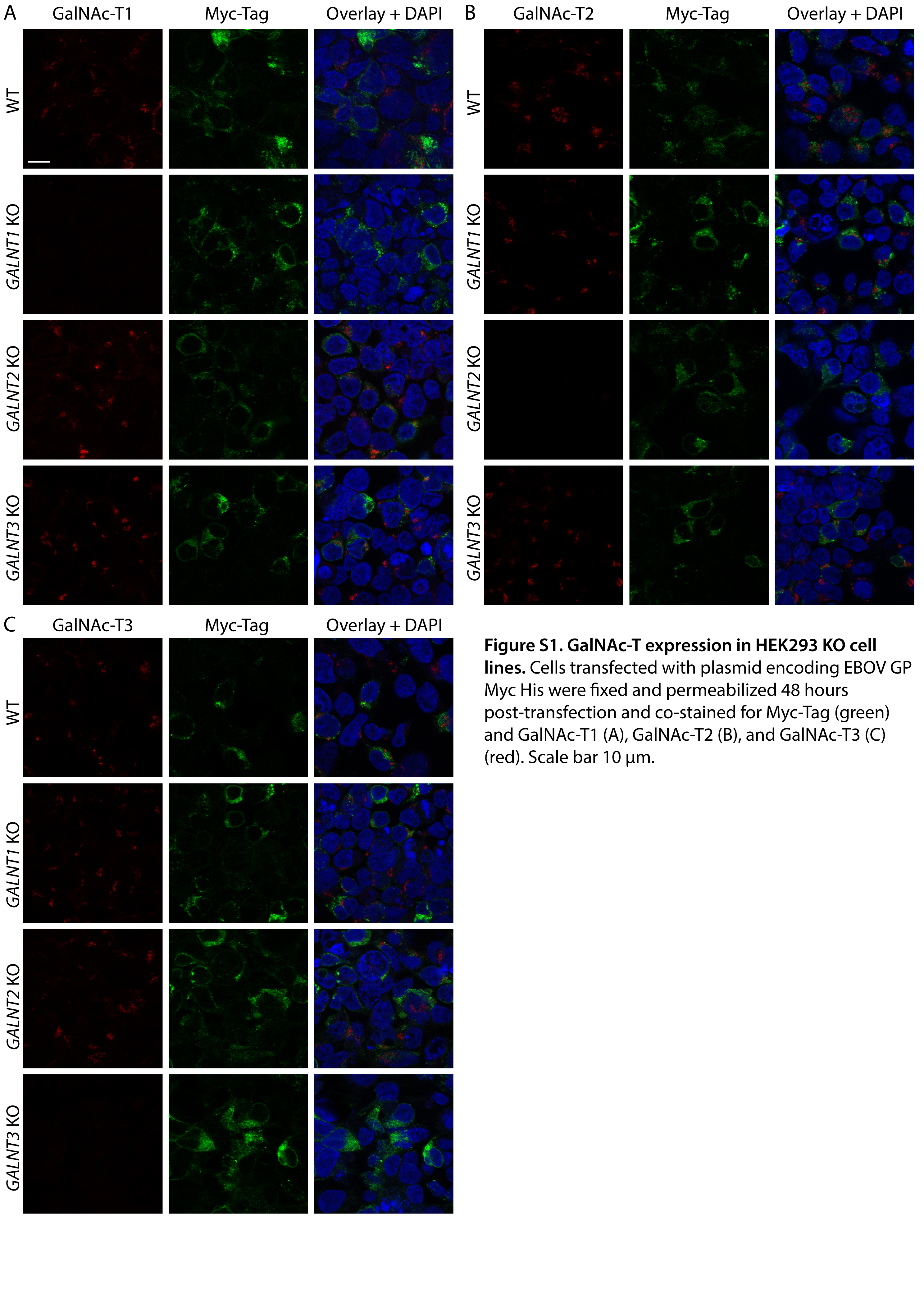

### Figure S2

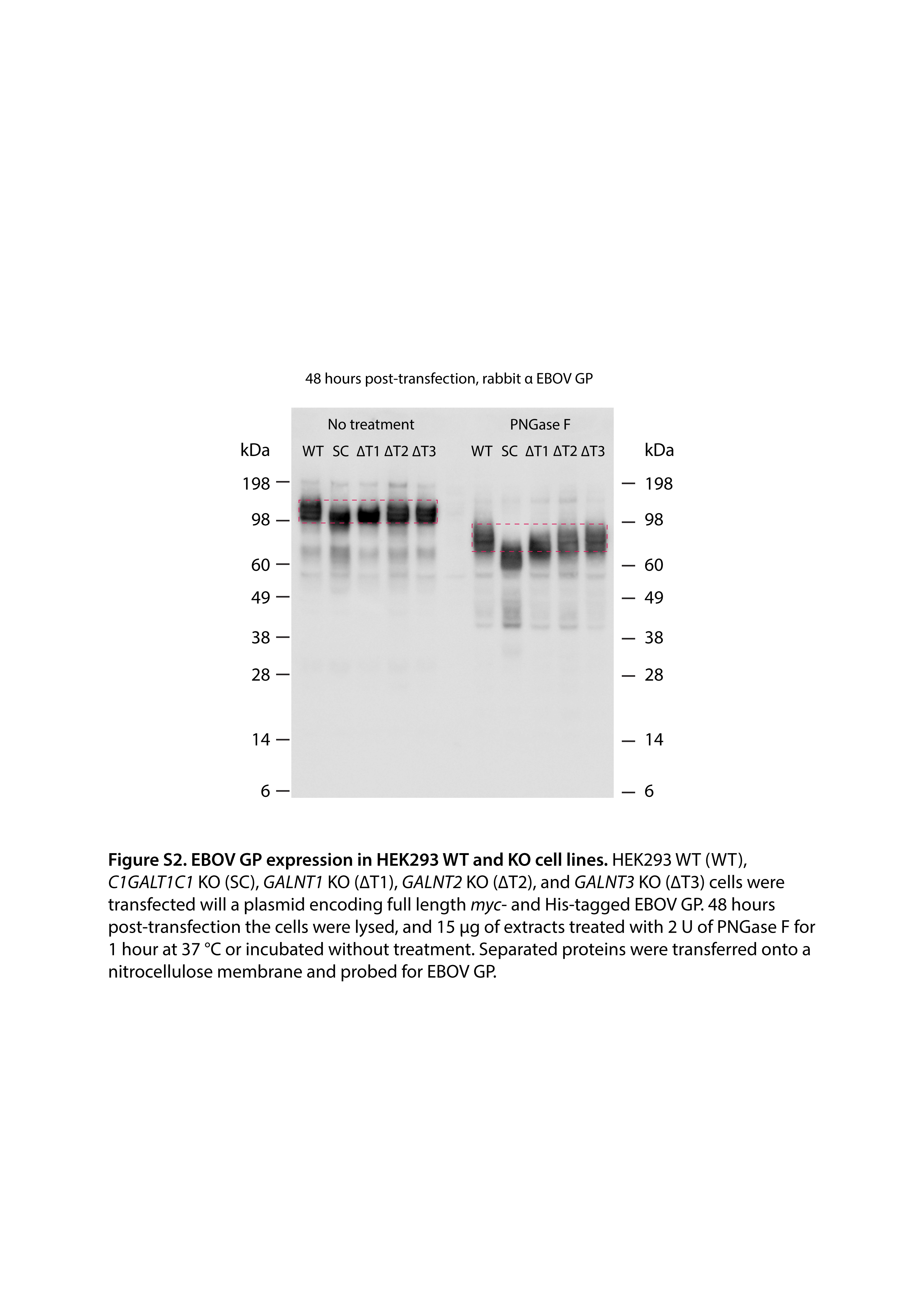

### Figure S3

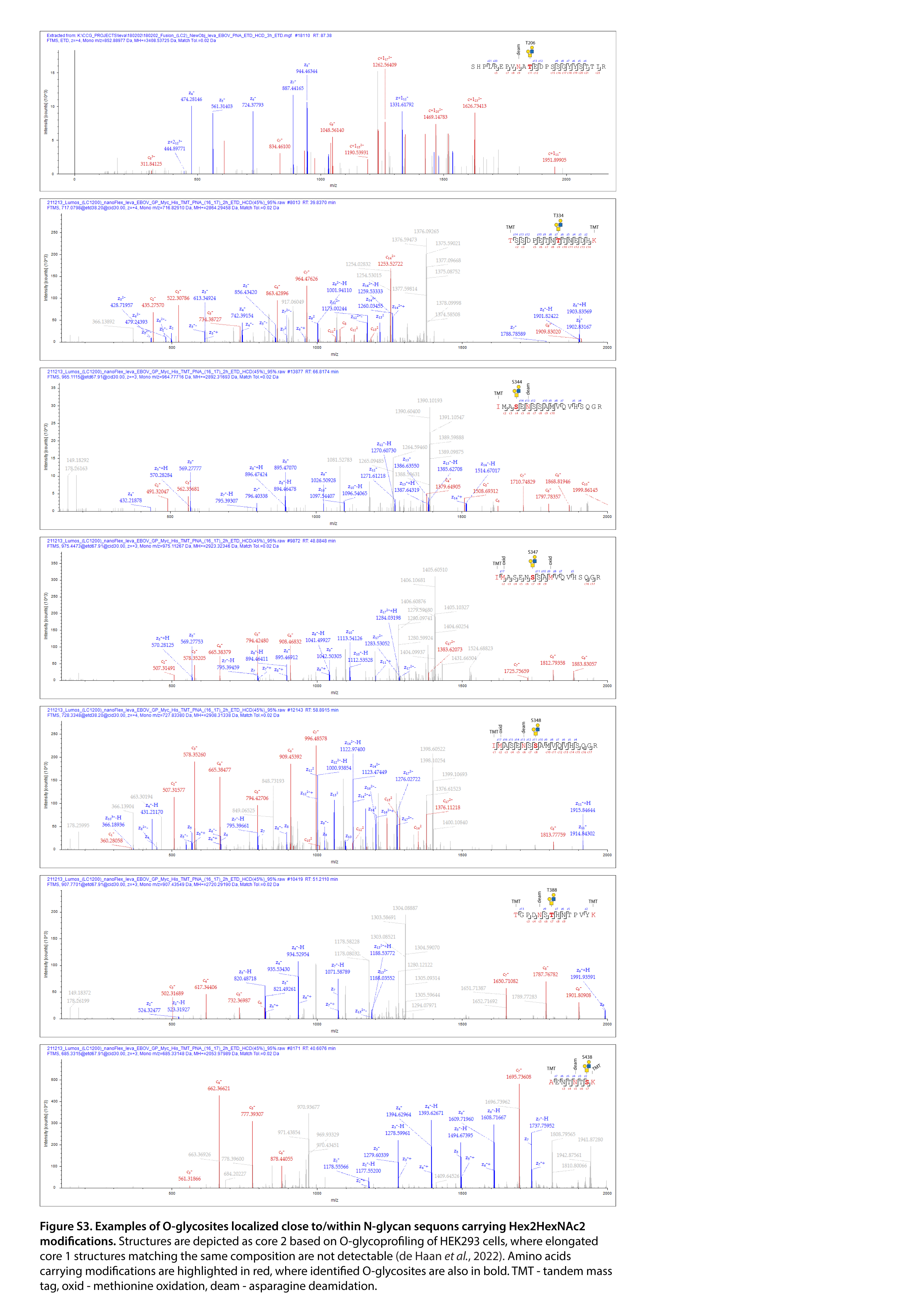
